# Circular RNA RORβ regulates TGFBR1 by decoying miR-140 in alcohol-exposed lungs and fibroblasts

**DOI:** 10.1101/2022.05.27.492519

**Authors:** Viranuj Sueblinvong, Xian Fan, Raven Williams, Bum-Yong Kang

## Abstract

Alcohol ingestion exaggerates transforming growth factor-beta 1 (TGFβ1) expression and signaling leading to fibroproliferation. Inhibition of TGFβ receptor type 1 (TGFβR1) mitigates the effect of TGFβ1 signaling. We showed that alcohol can modulate microRNA (miRNA) expressions. The mechanism by which alcohol modulates microRNA and how it ties to TGFβ1 signaling has not been well elucidated. Circular RNA (circRNAs or circ) emerges as a potential therapeutic target based on its stability, tissue specificity, and its ability to modify miRNAs. In this study, we showed that alcohol upregulates TGFβR1 and circRNA form of retinoic acid receptor-related orphan receptor beta (circ-RORβ) in lung fibroblasts (LF) and the lung. We identified miR-140 to have binding sites for both TGFβR1 3’ UTR and circ-RORβ and alcohol attenuated miR-140 expression in LF and the lung. We demonstrated that inhibition of circ-RORβ upregulated miR-140 and completely abrogated alcohol-induced miR-140 suppression. We further showed that inhibition of circ-RORβ attenuated alcohol-induced TGFβR1, fibronectin (FN1), and α-smooth muscle actin (αSMA) expressions and myofibroblast development as seen by an attenuation of αSMA stress fiber formation in LF. Taken together, these findings identify circ-RORβ-miR-140-TGFβR1 axis as a novel mechanism by which alcohol induces TGFβ1 signaling and promotes FMD.

**Highlights:** Alcohol induces circ-RORβ expression in lung fibroblasts

Circ-RORβ regulates TGFβR1 by decoying miR-140 in lung fibroblasts

Inhibition of Circ-RORβ restores miR-140 expression

Inhibition of Circ-RORβ mitigates alcohol-mediated myofibroblast differentiation

This is the first description of circ-RORβ functional significance in lung fibroblast

## Introduction

It has been well-recognized that chronic alcohol abuse independently increases the incidence of and mortality from the acute respiratory distress syndrome (ARDS), a severe form of lung injury caused by diverse inflammatory stresses [1–3]. We demonstrated that alcohol exaggerated production and actions of TGFβ1 as a mechanism underlies fibroproliferative disrepair following lung injury in the experimental model [4–6]. We further showed that inhibition of TGFβ1 receptor type 1 (TGFβR1) kinase (also known as activin-like kinase 5 or ALK5) mitigates the effect of alcohol on lung fibroblasts [5, 6]. Taken together, these data suggested that TGFβR1 is one of the key molecular targets in alcoholic lung phenotype. However key mechanism(s) by which alcohol activates TGFβ1-TGFβR1 axis leading to increase lung susceptibility to injury and disrepair remains elusive.

Several miRNAs were shown to regulate TGFβ1 and its signaling molecules including miR-140 which were shown to target TGFβR1 [7–9]. Specifically, loss of miR-140 is associated with epithelial-to-mesenchymal transformation, myofibroblast differentiation, and fibrosis in several tissues including the liver and the lung [10, 11]. Further, it has been shown that there is a conserve pattern of miR-140 expression in the brain of alcoholic mouse and human [12]. However, less is known about miR-140 regulation in alcohol-exposed lung disease.

Retinoid acid-related orphan receptor (ROR) family consists of three members of the nuclear receptors (i.e. RORα, β, and γ) that are known to have ability to regulate anti-oxidative stress functions, and tissue repair and can function as transcription factor.[13] However, little is known about the expression and function of ROR family in alcohol-induced lung diseases.

Circular RNAs (circRNAs) is a class of lncRNA by which single-stranded RNA that are generated from the back-splicing of exons, introns, or both, 3’ and 5’ then form a covalent bond resulting in a close loop structure which are resistant to degradation by RNA exonucleases [14, 15]. They can be found across species and are evolutionally conserved [14]. The most well-recognized functional of circRNAs is its ability to regulate gene expression by abrogating (‘sponging’) targeted miRNAs [15]. Given their resistant to RNA exonucleases making them more stable compared to linear RNA, their specificity in its expression in pathological conditions, and their tissue-specific nature, circRNA have enormous potential as disease biomarkers and therapeutic targets [16]. We sought to identify the circular RNA which participates in regulatory mechanism of miR-140 in the lung and fibroblasts.

In this study, circRNA form of RORβ (circ-RORβ) was observed to be upregulated in alcohol-treated human and murine lung fibroblasts and murine lungs. Further, we identified a potential interaction between circ-RORβ, miR-140, and TGFβR1. We speculated that alcohol induces TGFβ1 signaling through modifying circ-RORβ-miR-140-TGFβR1 interactome leading to enhancement of fibroblast-to-myofibroblast differentiation (FMD) in alcohol-exposed experimental model and ultimately, priming the lung toward fibrotic disrepair following acute lung injury.

## Methods

### Animals and chronic alcohol ingestion

Three-month old C57BL/6J wild type mice were obtained from Jackson Laboratories (Bar Harbor, ME). Mice were fed without or with alcohol (20% v/v in drinking water) for 8 weeks as previously described,[4] and then lungs were collected for gene expression analysis and primary lung fibroblast (LF) isolation as described below. All studies were approved by the Institutional Animal Care Use and Committee (IACUC) at Emory University and conformed to institutional and Association for Assessment and Accreditation of Laboratory Animal Care International (AAALAC) standards of medical sciences’ ethical principles and guidelines for the humane treatment of laboratory animals.

### Cell culture and treatment

Mouse lung fibroblasts (MLF) were isolated from the lungs of three-month old C57BL/6J mice (Jackson Laboratories, Bar Harbor, ME) as previously described [17]. Cells were cultured and expanded in the complete MLF culture media (Dulbecco’s modified Eagle’s medium (DMEM)(Cellgro, Manassas, VA), 4.5 g/L glucose supplemented with 20% fetal bovine serum (FBS), 100 U/ml penicillin, and 100 U/ml streptomycin). MLF were used in experiments between 3-8 passages. Cells were subjected to alcohol treatment (60 mM)[17] while in DMEM culture media supplement with 5% FBS and 100 U/ml penicillin/streptomycin in closed tissue culture chamber with 60 mM alcohol in the bath within tissue culture incubator. Normal human lung fibroblasts (HLF) were obtained from ATCC (CCL-153) and were cultured in F-12K medium supplemented with 10% FBS per manufacturer protocol. Cells were subjected to alcohol treatment (60 mM, kept in alcohol chamber as described before) and ASO circ-RORβ (50 nM) as described below.

### In silico analysis for miRNA in correlation with circ-RORβ-TGFBR1 axis

To explore the potential role of circ-RORβ in alcohol-induced TGFβR1, we performed *in silico* analyses which showed that both circ-RORβ and TGFBR1 3’-UTR contain 56 common miRNA binding sites using Circular RNA Interactome [18] and TargetScans [19].

### Inhibition of circ-RORβ

HLF were transfected with custom-designed anti-sense oligonucleotide (ASO) circ-RORβ (IDT, 50 nM) or scramble ASO using Lipofectamine 3000 (Thermo Fisher Scientific, Waltham, MA). Six hours after transfection, serum-free media was replaced with DMEM culture media supplemented with 5% FBS, and cells were incubated overnight. Cells were treated with alcohol (60 mM) 24 hours after transfection, and then continued incubation in culture media supplemented with 5% FBS for 48 or 96 hours for gene expression and immunofluorescent analyses.

### Total RNA isolation and expression analysis

Total RNA was isolated from mouse lungs, HLF, and MLF using a commercial RNA isolation kit according to the manufacturer’s protocol (Zymo Research, Irvine, CA)[20]. First-strand cDNA was synthesized, and quantitative PCR was performed with primers for 18s, 9s, TGFβRI, fibronectin, αSMA, and circ-RORβ using iQ SYBR Green Supermix (Bio-Rad, Hercules, CA). The level of target messenger RNA (mRNA) and circ-RORβ expression was normalized to 18s or 9s housekeeping gene levels and relative expression values were calculated as previously described [20].

### MicroRNA (miRNA) isolation and expression analysis

The mirVana miRNA isolation kit was used to extract and purify miRNAs according to the manufacturer’s protocol (Thermo Fisher Scientific, Waltham, MA). cDNA was synthesized from 250 ng of miRNA using the miScript II RT kit (Qiagen, Valencia, CA). Quantitative PCR (qPCR) was performed for RNU6B (Qiagen) and miR-140 (Invitrogen) expression using the QuantiTect SYBR Green PCR Kit. The level of target miR-140 expression was normalized to RNU6B. The relative levels were determined by the comparative cycle threshold method as previously described [7].

### Immunofluorescence

HLFs were cultured in 4-well chamber slides (Nunc Lab-Tek) ± Alcohol (60 mM) ± ASO circ-RORβ (50 nM) for 96 hours. Cells were fixed, permeabilized, and then stained for αSMA overnight using rabbit anti-human αSMA antibody at a 1:500 dilution at 4°C (Abcam, San Francisco, CA). The chamber slides were incubated with a goat anti-rabbit secondary antibody (Invitrogen, Carlsbad, CA) and DAPI (Vector Labs, Burlingame, CA) and examined using an Olympus BX-41 fluorescence microscope (Olympus, Center Valley, PA). Cell count was performed from four random fields per sample (x20 magnification) from triplicate experiments. Data were expressed as percent of cells with αSMA stress fiber formation compared with total number of cells.

### Statistical analyses

Unpaired two tailed t-tests or one-way analysis of variance (ANOVA) was utilized for comparisons between groups using GraphPad Prism and GraphPad InStat version 7 (San Diego, CA). If statistical significance was reached following one-way ANOVA, post-hoc analysis using Dunnett’s correction method was performed. Data are expressed as means and standard errors. Significant differences were accepted at a *p* value of <0.05.

## Results

### Alcohol increases TGFβR1 expression in lung fibroblasts and lungs

To examine whether alcohol regulates TGFβR1 expression in lung fibroblasts and lungs, human and mouse LFs were treated with alcohol for 48 hours in vitro, and mice were fed for 8 weeks. As shown in Fig. 1, alcohol significantly induced TGFβR1 expression in HLF compared to control condition (**Fig.1A**, p<0.05). Furthermore, to collaborate with our previous findings, we assessed TGFβR1 expression in alcohol-exposed murine models and showed that alcohol upregulated TGFβR1 expression in both the MLF and mouse lungs as similarly shown in HLF (**Fig.1B-C**, p<0.05).

**Figure 1.**
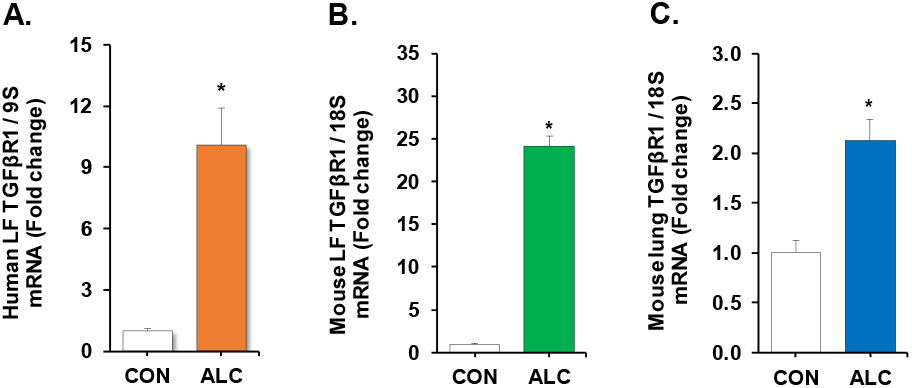
Alcohol induces TGFβ receptor type 1 expression in the lung and lung fibroblasts. (A) Human lung fibroblasts (HLF) were treated with alcohol (ALC, 60 mM, 48h). In parallel, (B) murine primary lung fibroblasts (MLF) were treated with alcohol (ALC, 60 mM, 48h). (C) Alcohol-fed mice were sacrificed after 8 weeks of alcohol ingestion (ALC; 20% v/v of ethanol in drinking water, ALC) along with their control-fed counterparts (CON). Samples were harvested and analyzed for TGFβ receptor type 1 (TGFβR1) expression by qPCR. N=4-7. *p<0.05.

### Alcohol induces circ-RORβ in the lung and lung fibroblasts

We sought to determine whether alcohol modulates circ-RORβ in alcohol-exposed lung fibroblasts and lung. As illustrated in Fig. 2, alcohol significantly induces circ-RORβ expression in HLF (**Fig.2A**, p<0.05). In parallel, we showed that alcohol induces circ-RORβ in MLF and mouse lungs similar to the alcohol-treated HLF (**Fig.2B-C**, p<0.05). To our knowledge, this is the first report of circ-RORβ in the literature.

**Figure 2.**
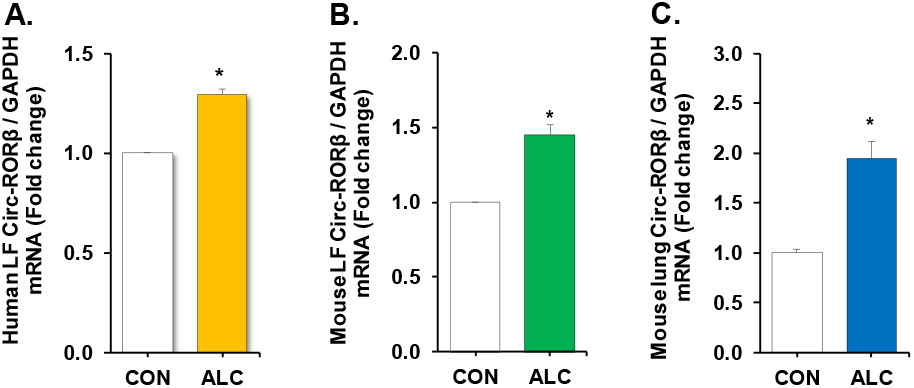
Alcohol induces circ-RORβ expression in the lung and lung fibroblasts. (A) Human lung fibroblasts (HLF) were treated with alcohol (ALC, 60 mM, 48h). In parallel, (B) murine primary lung fibroblasts (MLF) were treated with alcohol (ALC, 60 mM, 48h). (C) Alcohol-fed mice were sacrificed after 8 weeks of alcohol ingestion (ALC; 20% v/v of ethanol in drinking water, ALC) along with their control-fed counterparts (CON). Samples were harvested and analyzed for circ-RORβ expression by qPCR. N=4-6. *p<0.05.

### Circ-RORβ is identified as a competing endogenous RNA regulating miR-140-TGFβR1 axis

*In silico* analyses for miRNAs which can regulate TGFβR1 and interact with circ-RORβ by using Circular RNA Interactome and Target Scan 8.0, identified a total of 56 highly conserved miRNAs binding sites for human circ-RORβ and TGFβR1 3’UTR (**data not shown**). We analyzed the top 10 miRNAs expression with the highest target score as listed in **Fig.3A** and found that only miR-140 is significantly affected by alcohol treatment in HLF (**Fig.3B-C**, p<0.05). Specifically, alcohol exposure suppressed miR-140 expression in HLF (**Fig.3C**, p<0.05). Similarly, we also showed that alcohol significantly suppressed miR-140 expression in MLF and in mouse lungs as shown in **Fig. 3D-E** (p<0.05). These data suggest the correlation between alcohol and miR-140 in lung fibroblasts and the lung.

**Figure 3.**
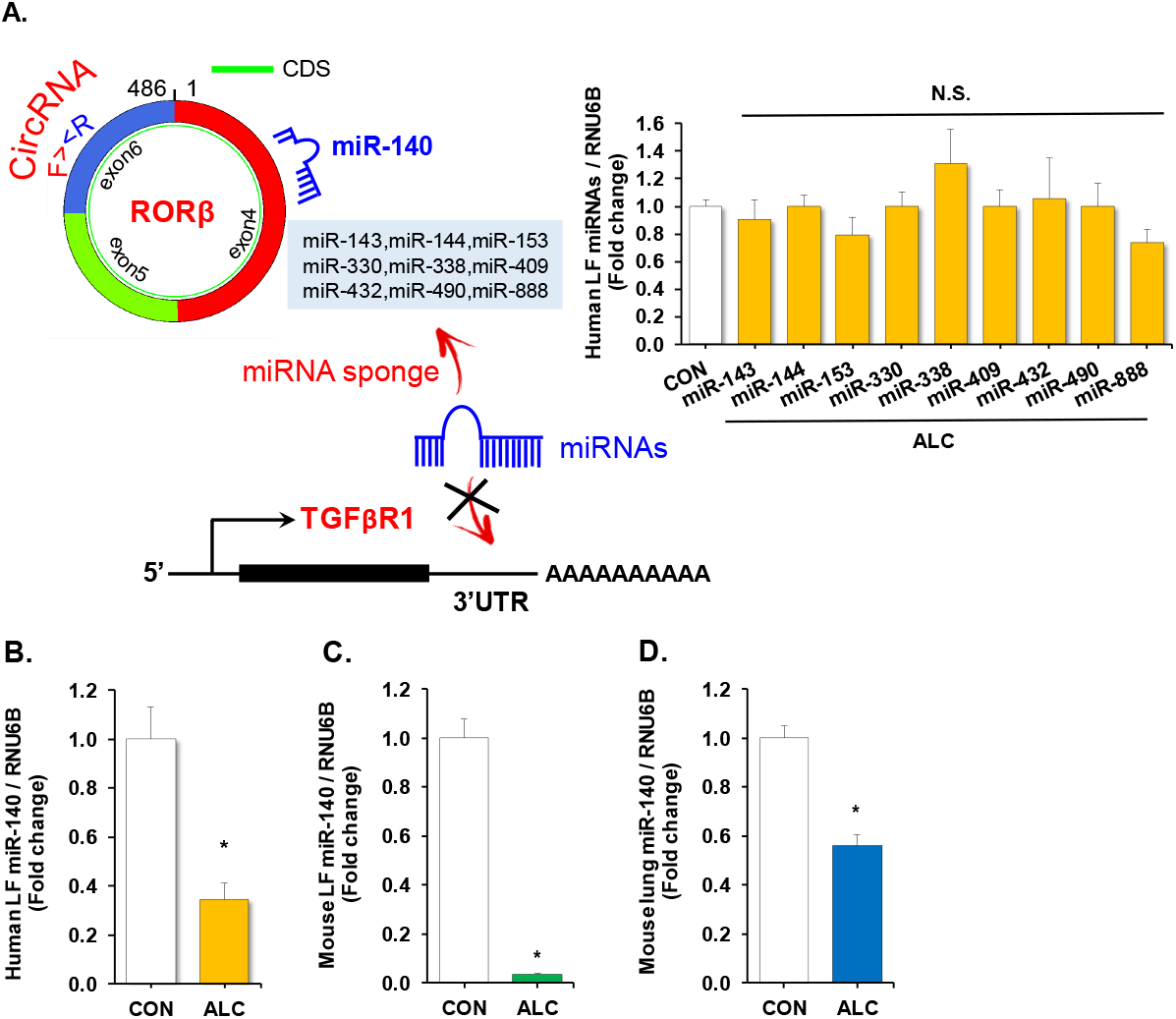
MiR-140 identified as a competing endogenous RNA for circ-RORβ and TGFβ receptor 1. (A) Image shows the top 10 miRNAs which could interact with both circ-RORβ and TGFβR1. (B) Human lung fibroblasts (HLF) were treated with alcohol (ALC, 60 mM, 48h), then analyzed for miR-143, miR-144, miR-153, miR-330, miR-330, miR-338, miR-409, miR-432, miR-490, and miR-888 by qPCR. (C) alcohol-treated HLF (ALC, 60 mM, 48h) were analyzed for miR-140 expression by qPCR. In parallel, (D) alcohol-treated murine primary lung fibroblasts (MLF) (ALC, 60 mM, 48h) and (E) the lung from alcohol-fed mice (ALC; 20% v/v of ethanol in drinking water, ALC) were analyzed for miR-140 expression by qPCR. N=3-7. *p<0.05.

### Inhibition of circ-RORβ attenuates alcohol-mediated miR-140 suppression and restores alcohol-induced fibroblast-to-myofibroblast differentiation

To assess the regulatory function of circ-RORβ on miR-140, circ-RORβ was inhibited in HLF using ASO. We showed that inhibition of circ-RORβ significantly upregulated miR-140 expression and abrogated alcohol-mediated miR-140 suppression in HLF (**Fig.4A**). To further determine the role of circ-RORβ in alcohol-induced lung fibroblast-to-myofibroblast differentiation (FMD), anti-sense oligonucleotides (ASO) against circ-RORβ was utilized. We showed that inhibition of circ-RORβ did not affect TGFβR1 expression in the absence of alcohol (**Fig.4B**). However, inhibition of circ-RORβ completely abolished alcohol-induced TGFβR1 expression in HLF (**Fig.4C**, p<0.05). We further assess the effect of circ-RORβ on FMD by analyzing the downstream genes to the TGFβR1 signaling including fibronectin and αSMA. There was no change in both fibronectin and αSMA expression in the absence of alcohol after blocking RORβ like its effect on TGFβR1 (**Fig.4C-D**). However, inhibition of circ-RORβ mitigated alcohol-induced fibronectin and αSMA expression in HLF (**Fig.4C-D**, p<0.05). In parallel, we assessed the function of circ-RORβ on FMD development by analyzing αSMA stress fibers formation. We showed that alcohol significantly induced FMD similar to the effect of TGFβ1 treatment (**Fig.4E**). Inhibition of circ-RORβ mitigated the numbers of FMD as seen by less cells with αSMA stress fibers compared to cells treated with TGFβ1 or alcohol. Taken together, these data demonstrate the functional significance of circ-RORβ in alcohol-induced FMD.

**Figure 4.**
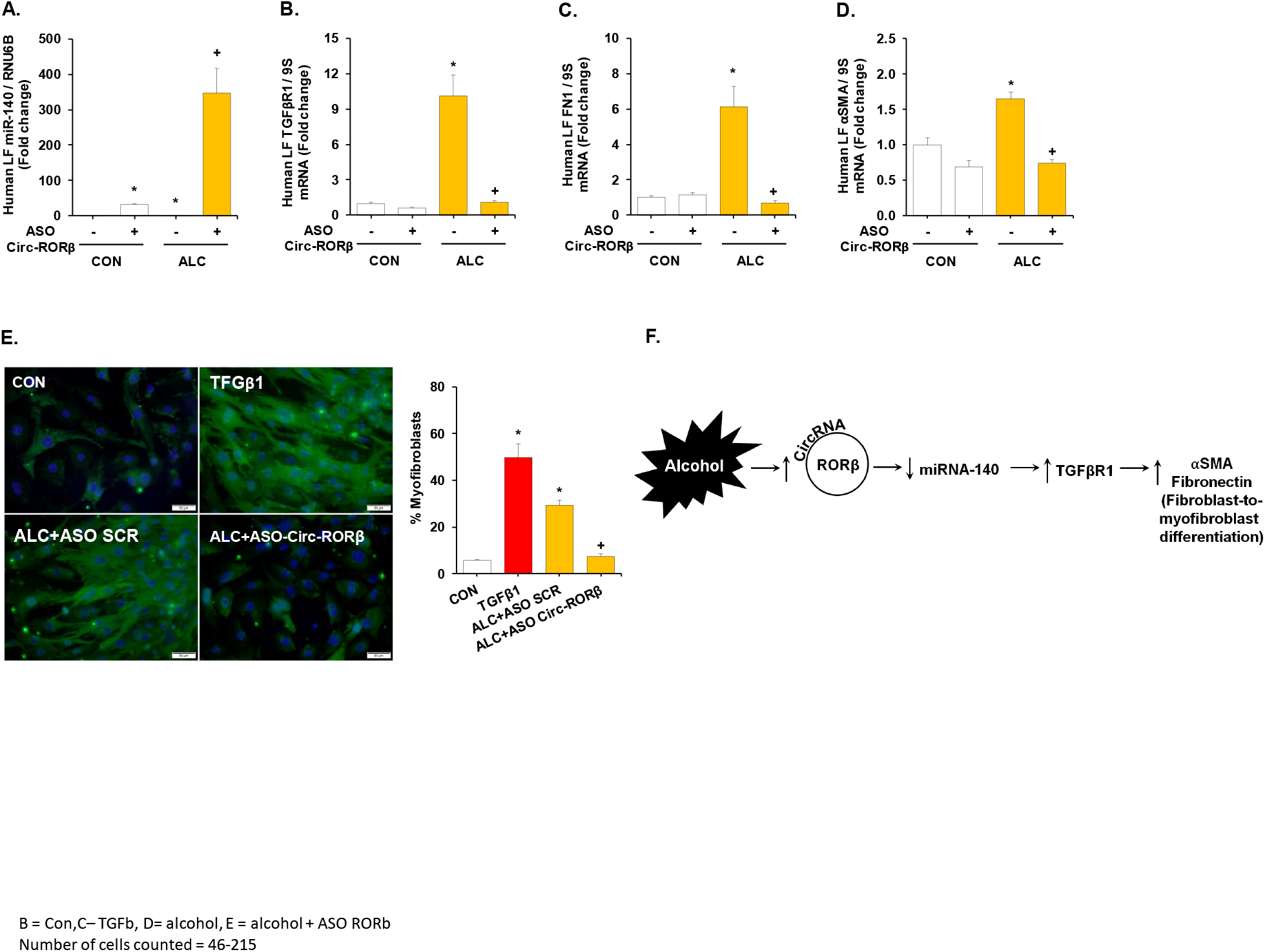
Inhibition of circ-RORβ attenuates alcohol-induced TGFβR1 and alcohol-induced fibroblast-to-myofibroblast differentiation (FMD). Human LFs (HLF) were transfected with synthetic anti-sense oligonucleotides against circ-RORβ (ASO circ-RORβ; 50 nM) or scrambled ASO (50 nM) using lipofectamine 3000. At 24 hours following transfection, cells were treated without (CON) or with alcohol (ALC; 60 mM) for an additional 48 hours then analyzed for (A) miR-140 expressions, (B) TGFβR1 and markers for FMD including (C) fibronectin, and (D) αSMA expression by qPCR. In parallel, cells were maintained in media with or without alcohol (60 mM) for an additional 96 hours following ASO circ-RORβ transfection, then fixed and stained with rabbit anti-human αSMA (green) and DAPI (blue). Cells treated with recombinant human TGFβ1 (5 ng/ml) were used as a positive control. Cells were counted from 3 random 20x field. Total cells counted of 46-215. (E) Graph depicts percent of cells with stress fiber formation to total cells. Representative slide from untreated (CON), TGFβ1, scramble AOS-transfected followed by alcohol-treated (ALC+ASO SCR), and ASO circ-RORβ-transfected followed by alcohol-treated (ALC+ASO Circ-RORβ) were shown, 40x magnification. *p<0.05 compared to untreated group. +p<0.05 compared to alcohol-treated group. (F) Diagram summarizes the findings presented in this study. In lung fibroblasts, alcohol induces circ-RORβ expression which, in turn, neutralizes miR-140. This interaction leads to an upregulation of TGFβR1 expression in alcohol-treated lung fibroblasts. These sequential events promote fibroblasts to differentiate into myofibroblasts which underlies the susceptibility to injury and disrepair in alcoholic lungs.

## Discussion

Chronic alcohol ingestion increases the risk of developing ARDS in the clinical setting [1]. In an experimental model, we showed that alcohol promoted fibroproliferative disrepair through induction of TGFβ1 expression and signaling in the lung [4]. Using TGFβR1 kinase (or ALK5) inhibitor to block TGFβ1 signaling mitigates alcohol-induced oxidative stress in lung fibroblasts [5, 6]. In this study, we aimed to identify a mechanism by which alcohol activates TGFβ1 signaling with the focus on TGFβR1 and its competing endogenous RNAs. We utilized both human and mouse experimental models to collaborate with our previous findings in rodent models as well as for future experiments. We demonstrated that alcohol upregulated TGFβR1 in lung fibroblasts and mouse lungs. We identified circ-RORβ and miR-140 as competing endogenous RNAs to TGFβR1 as reflected alcohol induced circ-RORβ while suppressed miR-140 expression. Furthermore, inhibition of circ-RORβ mitigated alcohol-induced TGFβR1 expression and its downstream targets including fibronectin and α-smooth muscle actin (αSMA). In parallel, circ-RORβ inhibition also attenuated alcohol-induced fibroblast-to-myofibroblast differentiation (FMD). Lastly, inhibition of circ-RORβ restored miR-140 expression in alcohol-treated human lung fibroblasts (HLF). Taken together, these results demonstrate that alcohol induces TGFβ1 signaling through mediation of circ-RORβ-miR-140-TGFβR1 interactome in lung fibroblasts (**Fig. 4F**).

TGFβ1 is an evolutionarily conserved growth factor which is known to participate in fibrogenesis in various tissues [21–25]. TGFβ1 is secreted and stored in a latent form by binding to latency associated peptide [26]. Upon its activation, TGFβ1 binds to two heteromeric complexes of its receptor TGFβ receptor type 1 and type 2 (TGFβR1 and TGFβR2) resulting in activation of TGFβR1 kinase or ALK5 leading to activation of through a canonical or a non-canonical signaling pathway [26]. Our work previously showed that inhibition of TGFβR1 kinase resulted in attenuation of the alcohol effect in lung fibroblasts and alveolar epithelial cells [5, 27]. With these information, we analyzed the effect of alcohol on TGFβR1 and showed that alcohol induces TGFβR1 expression in lung fibroblasts and mouse lung while alcohol did not affect TGFβR2 expression in our experimental model (data not shown). This data is similar to the work published by Zhou et al showing that alcohol upregulated TGFβR1 expression in mouse hippocampus [28]. However, the mechanism by which alcohol regulates TGFβR1 is to be elucidated.

Retinoid-related orphan receptors (RORs) are a family of nuclear factors consisting of three subtypes including RORα, β, and γ [29]. RORβ were predominantly studied in disease associated with central nervous system pathology including epilepsy [30]. Alcohol was shown to upregulate RORβ in cerebral organoid generated from human induced pluripotent stem cells [31]. Very little data shows the importance of RORβ in the lung or how it might participate in tissue fibrogenesis. However, there are several published data linking long-noncoding ROR (lnc-ROR) to epithelial-mesenchymal transition and tissue fibrosis [32, 33]. Accordingly, we speculated that lnc-ROR might play a role in alcohol-induced lung susceptibility to injury and disrepair. We performed computer simulation analyses and identified RORβ gene that could form a circular form of a non-coding RNA or circRNA.

CircRNA is a type of lnc-RNA that emerges as a potential novel therapeutic target given its resistance to endonuclease and specificity to tissue and disease stage [14, 15]. CircRNA also was shown to have the ability to regulate gene splicing, parental gene transcription, interacting with target proteins, and influence protein translation [34]. However, the most well-recognized function of circRNA is its ability to sponge or neutralize miRNA resulting in a downregulation of target miRNA [15]. In this study, we assessed the effect of alcohol on circ-RORβ and show that alcohol upregulates circ-RORβ expression in lung fibroblasts and mouse lungs. To our knowledge, this is the very first report of circ-RORβ in literature. We next sought to identify a mechanism by which alcohol modulates TGFβR1 and how it is linked to circ-RORβ.

Several miRNAs were shown to participate in regulation of TGFβ1 and its signaling molecules including TGFβR1. We speculate that circ-RORβ mediated TGFβR1 through modulation of miRNA targeting TGFβR1. Through computer simulation analyses, we identified several miRNAs which could interact with both circ-RORβ and TGFβR1. Out of the top 10 miRNAs, alcohol affected miR-140 expression, but not other miRNA screened. These data suggest that miR-140 might be involved in alcohol-induced TGFβR1 in lung fibroblasts which is consistent with previous reports showing miR-140 involvement in fibrogenesis through its direct target of TGFβR1 in several experimental models [9, 10, 35]. We further confirmed the role of circ-RORβ in alcohol-induced FMD and showed that inhibition of circ-RORβ attenuated alcohol-induced TGFβR1 and FMD markers including fibronectin and αSMA. Most importantly, inhibition of circ-RORβ mitigated alcohol-induced myofibroblast differentiation as shown by the presence of αSMA stress fiber formation. Lastly, we showed that inhibition of circ-RORβ resulted in restoration of miR-140 in alcohol-treated lung fibroblasts. To our knowledge, this is the first report on circ-RORβ and linking circ-RORβ with miR-140-TGFβR1 axis.

Taken together, we present new evidence that alcohol mediates circ-RORβ-miR-140 axis enhances TGFβ1 signaling though upregulation of TGFβR1 in lung fibroblasts. More importantly, we showed that circ-RORβ might be involved in FMD which, in turn, leads to tissue fibrosis. Future studies to confirm our findings in the clinical setting are warranted. Therapies aiming to neutralize circ-RORβ could prevent the development of ARDS, reduce severity, and promote optimal lung recovery from ARDS in high-risk individuals.

## Ethics approval

All studies were approved by the IACUC at Emory University and conformed to institutional standards and the Swiss Academy of Medical Sciences’ ethical principles and guidelines for the humane treatment of laboratory animals.

## Authors’ contributions

BYK and VS designed the research project, BYK, VS, and XF performed experiments, analyzed data, interpreted the results of the experiments; BYK, HKQ, and VS drafted, edited, and revised the manuscript.

